# Genome Complexity Browser: estimation and visualization of prokaryote genome variability

**DOI:** 10.1101/713123

**Authors:** Alexander I Manolov, Dmitry N Konanov, Dmitry E Fedorov, Ivan S Osmolovsky, Elena N Ilina

**Affiliations:** Federal Research and Clinical Centre of Physical and Chemical Medicine, Federal Medical and Biological Agency of Russia, Moscow, Russian Federation

## Abstract

**Motivation:** Comparative genomics studies may be used to acquire new knowledge about chromosomal architecture - the rules to combine a set of genes in a genome of a living organism. Hundreds of thousands of prokaryote genomes were sequenced and assembled. Still, there is a lack of computational tools able to compare hundreds of genomes simultaneously, i.e. to find hotspots of genome rearrangements and horizontal gene transfer or to analyze which part of an operon is conservative and which is variable.

**Results:** We developed Genomic Complexity Browser (GCB), a tool that allows to visualize gene contexts in a graph form and evaluate genome variability of different parts of a prokaryotic chromosome. We introduce a measure called complexity, which is an indirect measure of genome variability. Intraspecies and interspecies comparisons reveal that regions with high complexity tend to be located in a similar context in different strains and species. While many of such hot spots are associated with prophages and pathogenicity islands, some of them lack these determinants and mechanisms that govern their dynamics are to be elucidated.

**Availability:** GCB is freely available as a web server at http://gcb.rcpcm.org and as a stand-alone application at https://github.com/DNKonanov/GCB

## 1 Introduction

Although highly variable, prokaryotic genomes do not represent simply a set of genes, they possess regularities, which are collectively termed “genome architecture” (Rocha, 2008; Touchon and Rocha, 2016; Hendrickson *et al.*, 2018). Rules of genome architecture can shed light on still unknown molecular mechanisms governing prokaryotic cell life and may be essential in the engineering of synthetic organisms.

One of the earliest experimental observations dealing with genome architecture of prokaryotes was made by A. Segall (Segall *et al.*, 1988) who experimentally introduced inversions in *Salmonella enterica* genome. While many of such inversions were neutral for bacteria fitness, some had a detrimental effect. It was suggested that the effect was due to the importance of correct location and orientation of replication termination sites (Sharma and Hill, 1995; Mahan and Roth, 1991).

Non-random localization of different genes may be important due to several factors. Genes located near the replication origin have higher copy numbers in fast-dividing cells – a so-called replication-associated gene dosage effect (Couturier and Rocha, 2006; Slager and Veening, 2016). Folding of the chromosome makes genes located in different parts of the chromosome close to each other in 3D space, which can be beneficial for the gene coding for a regulator and its targets (Dorman, 2013; Fritsche *et al.*, 2011). Effects of global regulators (such as H-NS) on gene expression were observed to depend on the location of the target gene (Brambilla and Sclavi, 2015) and transcriptional propensity also varies depending on chromosome position (Scholz *et al.*, 2019). Cooperative effects between RNA polymerases (Kim *et al.*, 2018) or supercoiling propagation effects may play a role in the transcriptional regulation of neighboring genes.

It was observed that horizontal gene transfer (HGT) events are preferentially localized in hot spots - chromosomal loci in which changes are observed much more frequently than in other regions (Touchon *et al.*, 2009). This might indicate that, although disruptions in genome architecture may result in decreased fitness of an organism, there are some places in the chromosome where changes can be introduced without inducing negative effects.

Here we present GCB, a tool that allows to estimate local genome variability and to visualize gene rearrangements in user-defined sets of genomes. Local genome variability is estimated using graph representation of gene order (neighborhood) with a here introduced measure called complexity. Complexity profiles may be used to identify hotspots of horizontal gene transfer and a graph-based visualization available in GCB allows to analyze patterns of genome changes events.

## 2 Methods

### 2.1 Data acquisition

To construct a dataset for the web server, we downloaded genomes for 143 prokaryotic species that had more than 50 genomes available from the RefSeq database. For each species, if the number of complete genomes available was higher than 50, then only complete genomes were used. If the number of available genomes was higher than 100, then exactly 100 genomes were randomly selected for further analysis. The only exception was *Escherichia coli* extended genome set, which contained 327 complete genomes available as of November 2017. All downloaded genomes were reannotated with Prokka ver 1.11 (Seemann, 2014) to achieve uniformity. Genes were assigned to orthologous groups with OrthoFinder ver. 2.2.6 (Emms and Kelly, 2015). Python scripts which are a part of the GCB application were used to parse OrthoFinder output, calculate genome complexity through a chromosome and generate subgraphs around genome regions of interest.

### 2.2 Graph construction

The algorithm for graph construction is the following: each orthologous group represents a node, and two nodes are connected by a directed edge if the corresponding genes are located sequentially in at least one genome in a set. The weight of the edge is calculated as the number of genomes in which corresponding genes are adjacent (see Figure 1A and 1B). Graph objects and their methods are implemented in gene-graph-lib library for Python 3, more information can be found in the library documentation at https://github.com/DNKonanov/gene_graph_lib.

**Fig. 1.**
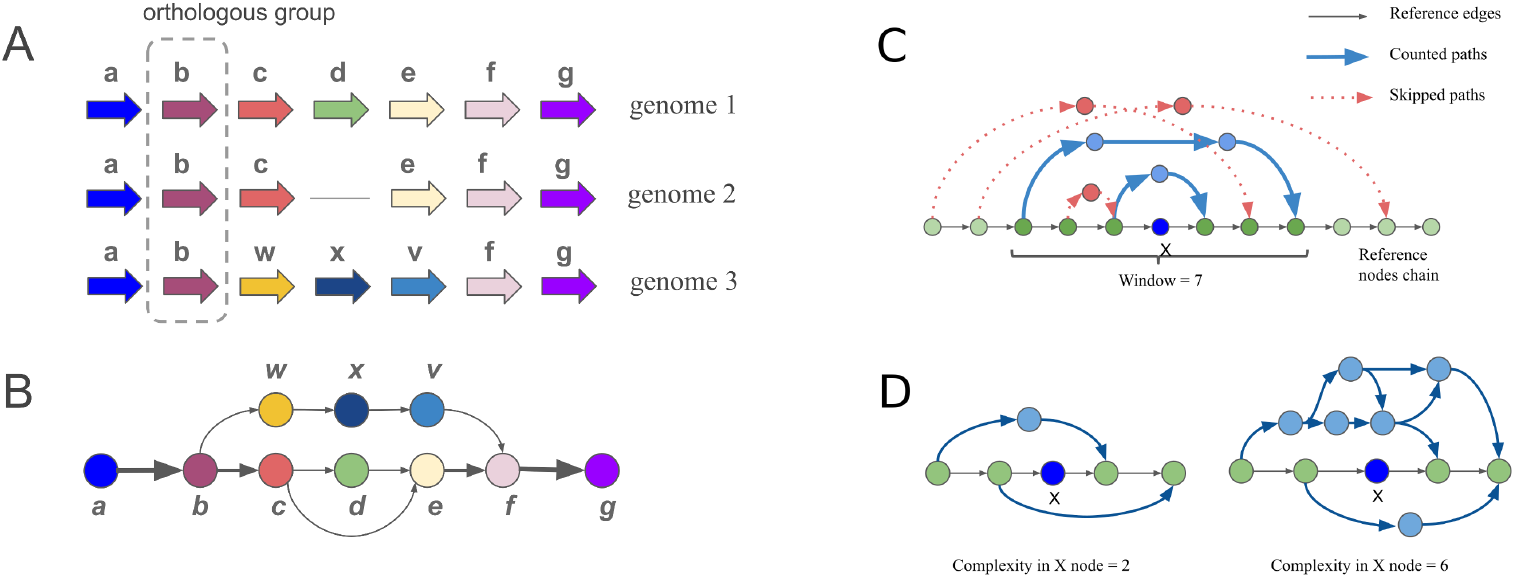
Principal scheme of a graph representation of gene order in a set of genomes and the complexity definition. To construct a graph, each orthology group is represented as a node. Nodes are connected by a directed edge if the corresponding genes are arranged sequentially in at least one genome in the set. A) genomes 1,2,3 represent three different hypothetical genomes. Arrows represent genes, genes from one orthologous group have the same color and letter designation. B) circles and arrows represent nodes and edges of the constructed graph. The weight of the edge (arrow width) is calculated as the number of genomes in which corresponding genes are located sequentially. C) Deviating paths for node X are defined as paths in the graph which bypass node X and are connected with an area of the reference node chain limited by the window parameter. D) Complexity value is defined by the number of paths deviating from reference node chain. There are two examples of calculating the deviating paths number, X is the considered node, deviating paths are shown with blue lines.

Because GCB uses directed graph representation of gene order, all genomes in a set should first be co-aligned in order to achieve the same orientation throughout the set. The algorithm for this step is listed in Supplementary Listings 1). Paralogous genes (orthologous groups which have more than one representative in one genome) are skipped by default but can be “orthologized” and added to the graph with an algorithm listed in Supplementary Listings 2 and described in Supplementary Notes 1.

#### Algorithm 1: COMPUTE COMPLEXITY

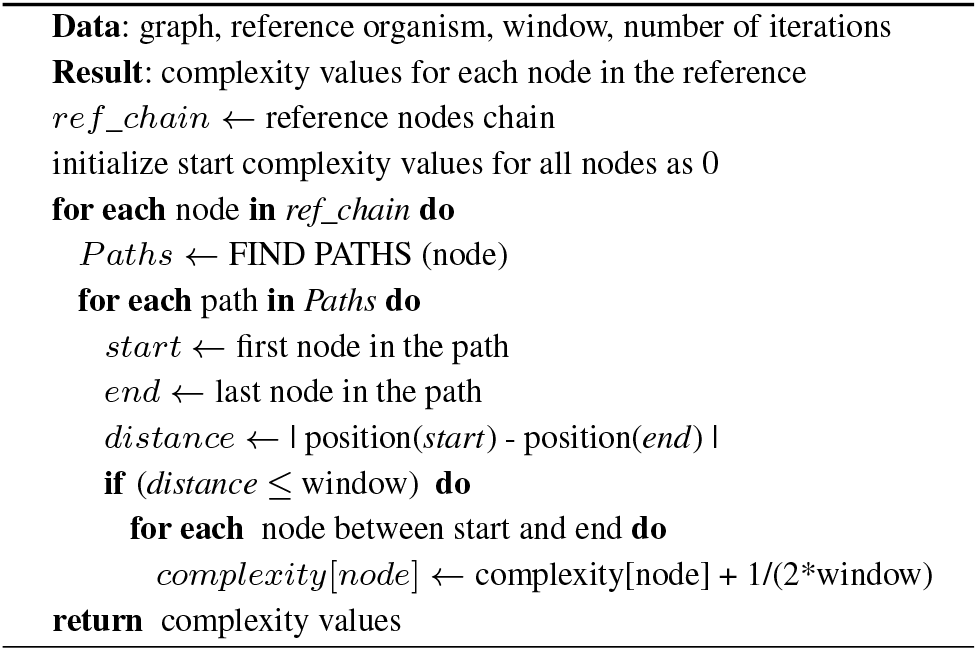

### 2.3 Genome complexity definition

Complexity values are calculated against one genome from the set that can be selected arbitrarily. This genome is extracted from the graph as a simple chain of nodes that is called the reference chain (Figure 1C). To calculate complexity in its node X, nodes from the reference chain in the range *±window/*2 around the node X are selected, and complexity is defined as the number of distinct paths in the graph that do not contain the node X but start and finish in the nodes from the selected range (deviating paths), see Figure 1C.

Complexity computing is an iterative algorithm that generates a set of possible deviating paths from each node in the reference genome (Algorithm 1). When a new unique deviating path is found, the algorithm adds 1/(2\**window*) value to all nodes in the reference between nodes coinciding with the start and the end of the deviating path.

The algorithm has the following user-defined parameters: *window* - the size of the area around node X to which deviating paths should be connected (default 20 nodes), *iterations* - number of random walk processes from each node (default 500)

### 2.4 Method validation

We supposed that calculated complexity in some region of a chromosome correlates with the probability of rearrangements in this region. To verify this assumption and validate the algorithm, a set of model genomes was generated by random rearrangements simulations. These simulations included random insertions, deletions, HGT events, and inversions. HGT and random insertion probabilities were chosen to be equal to deletion events probability to maintain genome length. The probability of inversion was chosen as 1/100 than others, in agreement with literature data that inversion events are less common than other types of rearrangements, such as deletions and duplication (Schmid and Roth, 1983). Localization of these changes in the chromosome depended on the user-defined distribution of rearrangements probability along the chromosome. Simulation algorithm is available in Supplementary Listings 5. Next, these model genomes were processed by the complexity computing algorithm and results were compared with input distributions (more details are available in Supplementary Notes 5).

### 2.5 Subgraph generation

To visualize a gene context in the region of interest, a subgraph representing this region is constructed. First, a subset of reference chain nodes representing the region of interest is added to the graph. Next, the algorithm iterates through other genomes from the set, and deviating paths limited to the selected region are added to the subgraph. If the length of the path is greater than the depth parameter, then the path is cropped, only start and end fragments (tails) of a fixed length (*tails* parameter, *tails* < *depth*) are added to the subgraph instead of the entire path. If the weight of some edge is less than the user-defined *minimal_edge_weight* parameter, this edge is not added to the subgraph. The subgraph generation algorithm is listed in Supplementary Listings 6).

### 2.6 Additional methods

Phylogenetic trees were inferred with Parsnp v1.2 (Treangen *et al.*, 2014). Retention indexes were calculated using RI function from R phangorn library (Schliep, 2010). To estimate similarity to the reference genome, all genomes were aligned with nucmer (Kurtz *et al.*, 2004) and similarity score was calculated as follows: all aligned reference genome ranges were reduced with IRanges R package (Lawrence *et al.*, 2013) and their total length was divided by reference genome length. All query genomes were sorted by this value and strains with the top 100 highest values were chosen. Nucmer was also used to detect synteny blocks in Figure 3 and Supplementary Figures 4-11. Prophages were detected with Phaster (Arndt *et al.*, 2016). Technical information (used libraries, frameworks, and their versions) is available in Supplementary Notes 6. SQLite database structure is described in Supplementary Notes 7. To obtain Figures 2A we used GCB with following parameters *tails* = 1, *minimal_edge* = 5. Figures 2B we used GCB with following parameters *tails* = 0, *minimal_edge* = 5. To obtain Figure 2C we used GCB with the following parameters: *window* = 20. Code to make Figures 2 and 3 is available at https://github.com/paraslonic/GCB_paper_code.

**Fig. 2.**
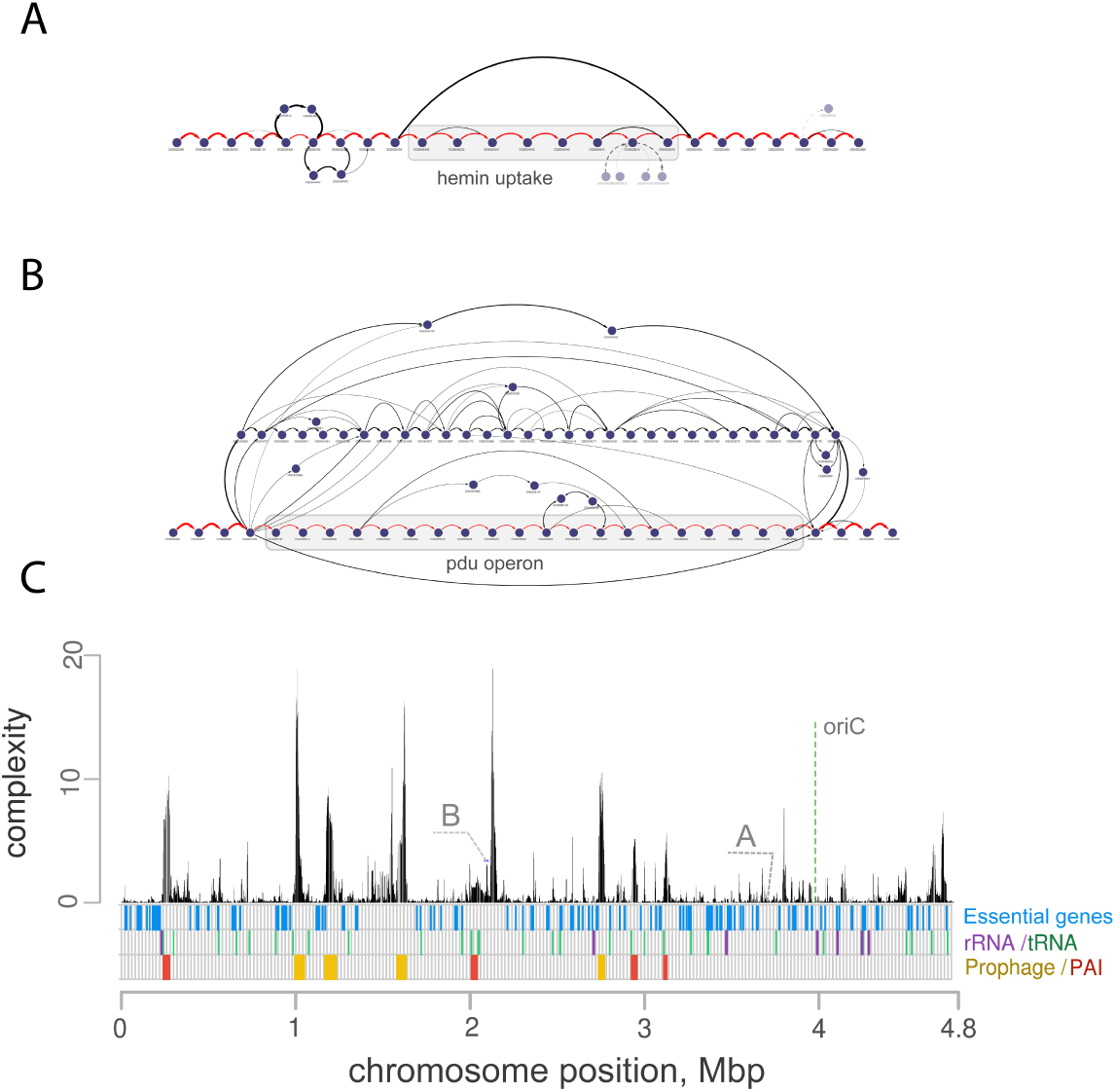
Regions with high complexity values are associated with prophages and pathogenicity islands. Subgraphs of the genomic regions in which A) hemin uptake and B) propanediol utilization operon are located. With dashed lines tails of long deviating paths are shown. The graph indicates that both operons are located in a conservative context. Genomes that do not contain hemin uptake operon do not contain other genes in the same context. In the case of propanediol utilization operon, in several genomes alternative and highly variable gene sets are present. Hmu operon is located on the 3691615-3700567 positions and pdu operon on the 2083448-2101340 positions of NCBI Reference Sequence NC_011993.1 (LF82 strain). C) Complexity profile for E. coli LF82 chromosome. Color bars at the bottom panel show the location of essential genes, ribosomal and transport RNA genes, regions with prophages and pathogenicity islands (PAI).

**Fig. 3.**
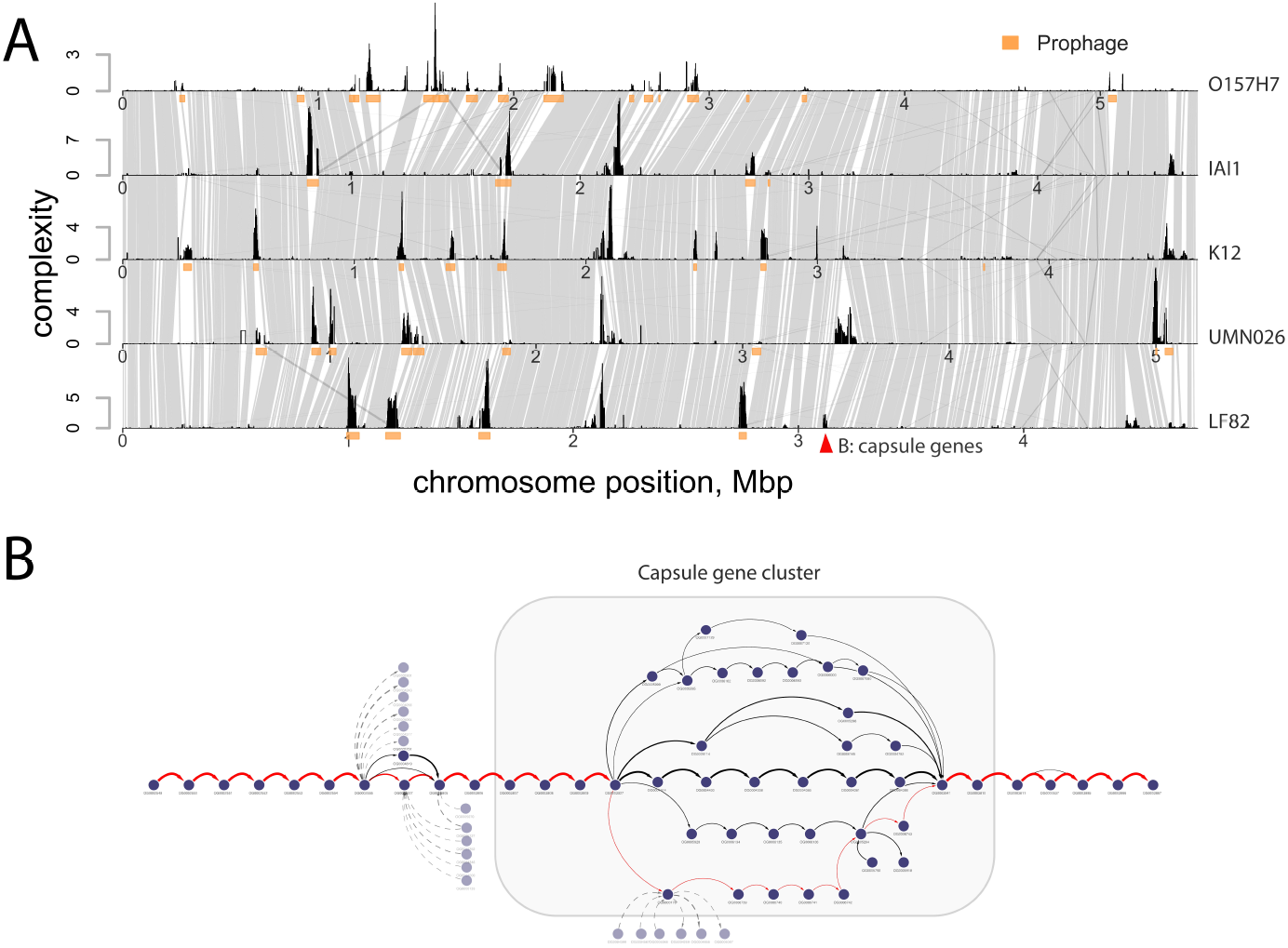
Regions with high complexity values are mainly located in a conservative context in different E. coli phylogroups. A) Genomes from five different phylogroups were used as references to compare complexity profiles. For each reference, 99 most similar genomes were selected and used to compute complexity values. Synteny is shown with gray blocks, orange boxes designate prophage regions. Red triangle shows the location of capsule genes cluster, which is located in the region of varying complexity (from low in IAI1 and O157H7 strains to moderate and relatively high in K12, LF82 and UMN026 strains). B) subgraph of capsule gene cluster shows the presence of conservative and variable parts of this operon in B2 clade (LF82 is reference genome). With dashed lines tails of long deviating paths are shown.

## 3 Results

### 3.1 Application

The GCB tool is available as a standalone application and as a web server. GCB web server uses precalculated data for 143 species and is available at http://gcb.rcpcm.org. The standalone browser-based tool and a set of scripts are available at https://github.com/DNKonanov/GCB.

GCB GUI consists of three main parts: 1) the top panel that allows selecting a genome and a region to work with, 2) the complexity plot showing complexity profile for selected genome and contig (in case if the assembly contains more than one contig), 3) the subgraph visualization form. Several settings are implemented to customize subgraph visualization to make it convenient for the analysis. Subgraph can be exported in JSON format and visualized with specialized software (i.e. Cytoscape) to prepare publication-ready images.

The server can also be run locally on a standard PC if the user needs to analyze a custom group of genomes. To estimate the genomic variability profile, the number of genomes should not be too small, few dozens or hundreds are typical values. The upper limit depends on the computational resources available to infer homology groups, which is the most computationally difficult step. Snakemake and Python scripts are provided to infer homology groups and to obtain a text file with complexity values or a database file, which can be imported to the local GCB server. Further details and instructions are provided in the user manual.

### 3.2 Subgraph visualization

Graph representation of gene order provides a convenient way to inspect visually the context of genes of interest and to identify conservative and variable gene combinations. GCB can construct and visualize subgraph - part of the graph containing the region of interest.

Figures 2 A and B show an example of subgraphs representing the gene context of the hemin uptake locus (hmu) and propanediol utilization operon (pdu) in 326 complete genomes of *Escherichia coli* from the RefSeq database (downloaded at the September 2017). The presence of *E. coli* harboring these operons in the intestinal microbiome was previously shown to be associated with Crohn’s disease (Dogan *et al.*, 2014; Viladomiu *et al.*, 2017; Rakitina *et al.*, 2017). While hmu operon is preferentially present in B2 phylogroup (Supplementary Figure 1, retention index = 1), pdu operon can be found in phylogenetically distinct strains of *E. coli*, and its presence is in low agreement with the phylogenetic tree (Supplementary Figure 2, retention index = 0.26). Visualization of subgraphs exported from GCB was made with Cytoscape (Shannon *et al.*, 2003), adherent-invasive *E. coli LF82* strain (Boudeau *et al.*, 1999) was used as a reference.

The graph visualization reveals that hmu operon Figure 2A is located in a conservative context, which means that the neighboring orthologous groups are the same in all strains in which it is present. The edge that bypasses the operon indicates that in some genomes the genes to the left and right of the operon are adjacent, which suggests that this operon was either deleted from ancestors of these genomes or inserted in other genomes. Graph visualization also indicates that one of the genes (hemin transport system permease, HmuU) or its close homologs are present in two alternative contexts.

Pdu operon (Figure 2B) is also located in a conservative context and some strains have no genes in that context. In this case, however, several genomes contain highly variable gene sets instead of pdu operon. These alternative sets include genes of iron transport (FepC, FcuA, HmuU), DNA mobilization (retroviral integrase core domain, transposase DDE Tnp ISL3), and others. Some variations in pdu operon itself are also visible and reflect different known operon variants (Rakitina *et al.*, 2017).

### 3.3 Complexity profiles

In a set of genomes with identical gene order, each node in the resulting graph will have two edges. Any gene rearrangements result in the addition of new edges. We hypothesized that the number of distinct paths in a subgraph representing a genomic region will monotonically depend on the frequency of gene rearrangements in this region. We implemented the algorithm (listed in Supplementary Listings 3 and described in Supplementary Notes 3) to count the number of distinct random walks in a subgraph representing a given region of the reference genome, the value which we further call complexity of the region. By selecting subregions with the sliding window we get the complexity profile of the reference genome. The width of the sliding window can be set by the user, the default value of 20 was used for the results described below.

To verify our approach, we performed a number of simulations in which we suggested that the probability of genomic rearrangement events (HGT, deletion, translocation) is non uniformly distributed along the chromosome. The algorithm is listed in Supplementary Listings 5 and described in Supplementary Notes 5. We used three patterns to generate profiles of such probabilities: sinusoidal, rectangle, and sawtooth and performed 10 independent simulations for each pattern. The results of our method were in good correspondence with the predefined distribution (R-square mean > 0.8, Spearman correlation > 0.7, FDR corrected p-value < 10^*−*300^), the comparisons of initial and inferred profiles are presented in Supplementary Notes 5.

Figure 2C shows the complexity profile of *E. coli LF82* chromosome inferred with our algorithm. As expected, the regions with a higher density of essential genes have lower complexity values. On the contrary, pathogenicity islands and prophages are associated with the regions with higher complexity. At the same time, there are chromosomal loci with relatively high complexity values which have no identifiable associates (no phage-like or transposon-associated genes). The comparison of the complexity profile with 3C experimental datasets and simulation-based chromosomal models did not reveal significant associations (Supplementary Notes 8).

The proposed method can also be used to compare variability profiles of different species and intraspecies structures. Figure 3A shows a comparison of complexity profiles for different *E. coli* phylogroups (Clermont *et al.*, 2013). For each of the five large phylogroups (A, B1, B2, D, E) we selected one reference strain and 99 most similar strains from 5466 RefSeq genomes (both finished and draft assemblies), see Supplementary Figures 3 for the resulting phylogenetic tree of the 500 selected genomes. Then complexity profiles for each reference genome were inferred. To compute complexity independently for each clade only genomes from the corresponding clade were used for each of the references. This comparison reveals that many of the rearrangement hot spots are conservative and located in the same context in the genomes of the strains belonging to different phylogroups. The majority of them contain prophages (denoted with an orange bar below the complexity profile), but some do not include phage-associated genes. Transient hotspots (with high complexity in some clades and low complexity in others) can also be observed.

We noticed that one of the transient hotspots near the pheV tRNA gene contains genes for capsule type II synthesis and export (red triangle on Figure 3A, chromosomal coordinates, and NCBI locus tags: CU651637:3111444-3128026, LF82_461-LF82_474). The visualization of this region in *LF82* genome from B2 clade (Figure 3B) reveals the presence of a highly variable region inside this operon, which was previously described (Johnson and O’Bryan, 2004). Tails of long paths which bypasses the operon are shown with dashed lines, they indicate that this operon is located inside the variable region of larger size.

The online tool contains precalculated complexity profiles for 141 bacterial and 2 archaeal species. Supplementary Figure 4 shows the comparison of 68 genomes for which exactly 100 genomes were used to compute complexity profiles and Supplementary Figure 5 for all available organisms. It can be seen that a number of species have similar complexity distributions through chromosome (Supplementary Figures 6-11).

## 4 Discussion

Hotspots of genome instability were described for a number of bacterial species. In (Oliveira *et al.*, 2017) the authors analyzed HGT hot spots for 80 bacterial species. They concluded that many hotspots lack mobile genetic elements and proposed that homologous recombination is mainly responsible for the variability of those loci. The factors that determine the location of hot spots, their emergence, and elimination, are still an open question. To our knowledge, GCB is the first tool that allows identifying genome rearrangement hot spots based on a user-defined set of genomes. This provides a way to study dynamics of hot spots, changes in their intensity and location on different levels ranging from intraspecies structures like phylogroups or ecotypes to interspecies and intergenus comparisons. While the dynamics of hot spots are primarily of fundamental interest, practical applications may also be expected in the field of synthetic biology, or in search for a new way to control the spread of antibiotic resistance.

We compared complexity profiles between different species and, in the case of *E. coli*, between different phylogroups. We observed that, as a rule, when genomes are close enough for the large synteny blocks to be detected (with blast or nucmer tool), then complexity profiles look similar: the regions with high complexity values are surrounded with low complexity regions forming the same conservative context in different groups of organisms. The analysis of complexity profiles of *E. coli* revealed that many hotspots are located in the prophage or pathogenicity islands integration sites, and site-specific mechanisms could govern their conservative location. Some hotspots lack such factors and reasons for their conservative location are still to be elucidated. We had hypothesized that chromosome folding may influence hotspots location, but no evidence of this was found by comparison of complexity profiles with 3C data and chromosomal folding simulations available from (Lioy *et al.*, 2018) and (Hacker *et al.*, 2017) correspondingly (Supplementary Notes 8).

While complexity values calculated by GCB reflect the intensity of gene rearrangement process, gene context visualization in a graph form reveals it details. One of the scenarios is the variation of alternative gene sets, and another is the process in which different changes overlap each other. While the first type of variability is in a good agreement with the mechanism proposed in (Oliveira *et al.*, 2017), the latter is not. The case of locus containing propanediol utilization operon (pdu) shows a combination of the two. The operon is present in a subset of phylogenetically distinct strains always in the same context, which means that it was either deleted from or inserted into some of the genomes. In either case, its gene content is rather stable with a relatively low number of observed variants. The opposite is true for the strains that do not contain this operon but have an alternative set of genes in its place. In this case, many overlapping changes are observed. It may be suggested that the reason for such difference lies in the functional importance (fitness benefits) of the operon for its carriers and “egoistic” or “parasitic” nature of alternative gene sets.

To our knowledge, gene neighborhood graph visualization is available only in FindMyFriends R package beside GCB. Our tool provides interactive opportunities and a number of visualization settings absent elsewhere.

The methods we proposed in this article are not universal, they are not suited for the detection of large genomic rearrangements (larger than window parameter, usually several dozens of genes) or changes in noncoding parts of the genome. Our methodology has also some drawbacks coming from its dependence on orthology inference accuracy. Here we used orthofinder tool (Emms and Kelly, 2015), which uses MCL graph clustering algorithm based on gene length normalized blast scores. We find this tool to be optimal in terms of efficiency and accuracy. On the other hand, it doesn’t take into account phylogenetic information and syntenic relationships between different genomes, and erroneous homology inference sometimes occur. Paralogous genes may fall into one group. In this case, the graph representation of the context becomes problematic. We implement two possible ways of dealing with paralogous genes in GCB: the default approach is to ignore them, the other is to perform artificial orthologization process (each paralogous gene with unique left and right context is denoted with a suffix and added to the graph). From our experience, the optimal strategy is to work in the default mode for explorative analysis and verify all conclusions in the paralogs orthologization mode. The graph layout process is also hard to automate. We use two layout algorithms (Dagre and Graphviz), but manual manipulations are often needed to make a clear layout, and Cytoscape (or other graph manipulation utils) is desirable to make publication-ready images.

Despite the above-mentioned drawbacks, we find the here proposed method of complexity analysis informative as it successfully identifies known rearrangement hot spots (prophages, integrons et al.), and we hope that GCB with its capacity of visualization and complexity assessment will find its application in the area of comparative genomics studies.

## Supporting information

Supplementary Listings

