## Supplementary Figures for "Genome Complexity Browser: estimation and visualization of prokaryote genome variability"

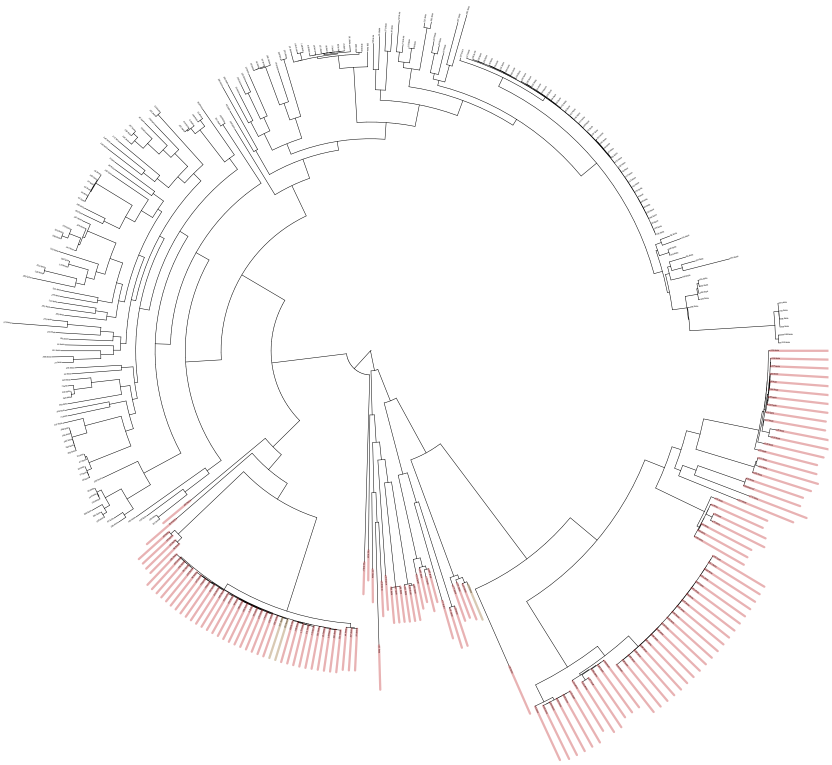

Figure 1: Hmu operon is in good correspondence with the phylogenetic tree of *E. coli*. Red bars denote genomes in which complete gene set from the operon is present, green bars denote genomes in which more than the half of operon genes are present.

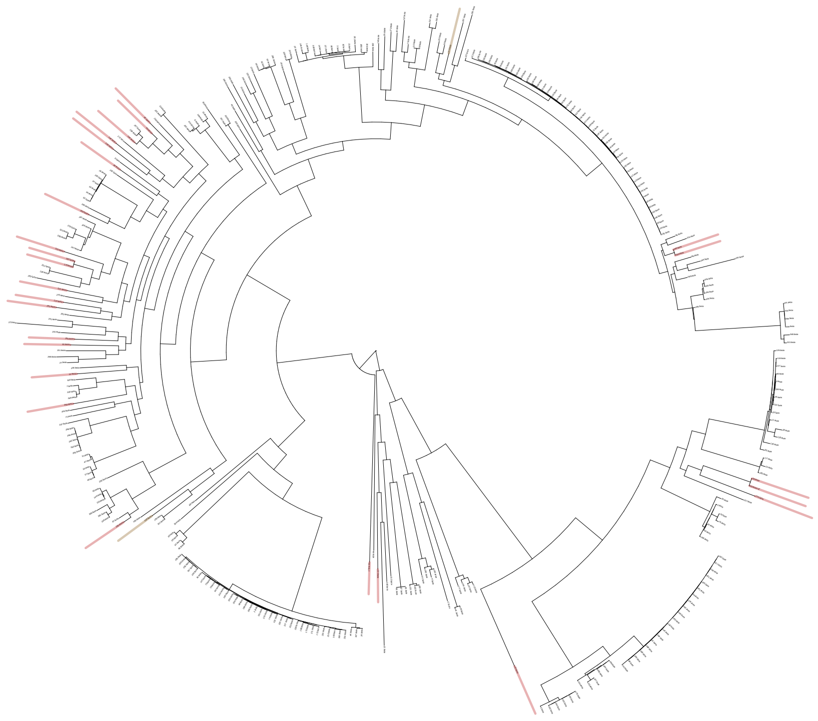

Figure 2: Pdu operon presence is poorly correlated with the phylogenetic tree of *E. coli*. Red bars denote genomes in which complete gene set from the operon is present, green bars denote genomes in which more than the half of operon genes are present.

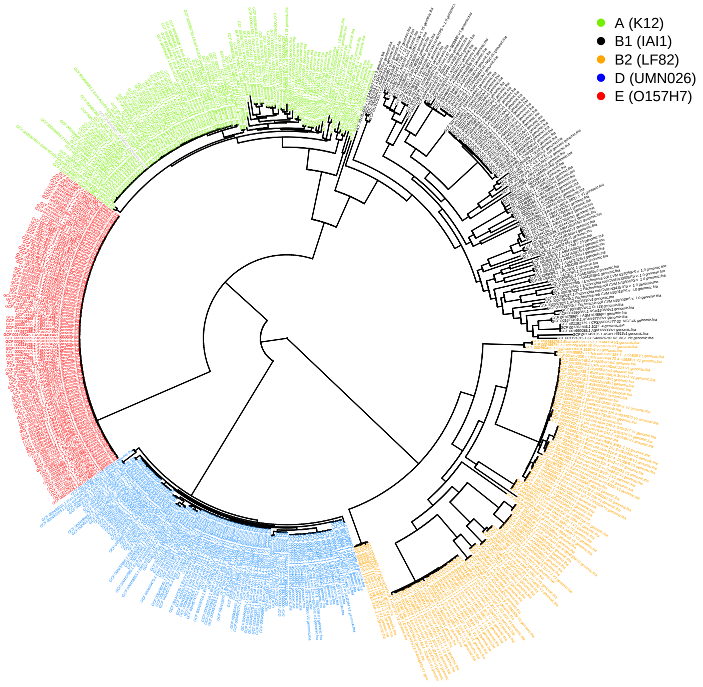


Figure 3: Phylogenetic tree of genomes selected to represent five *E. coli* phylogroups. First five genomes from each phylogroup were selected as references, then 99 genomes from RefSeq closest to each of them were added to the set giving 500 genomes total.

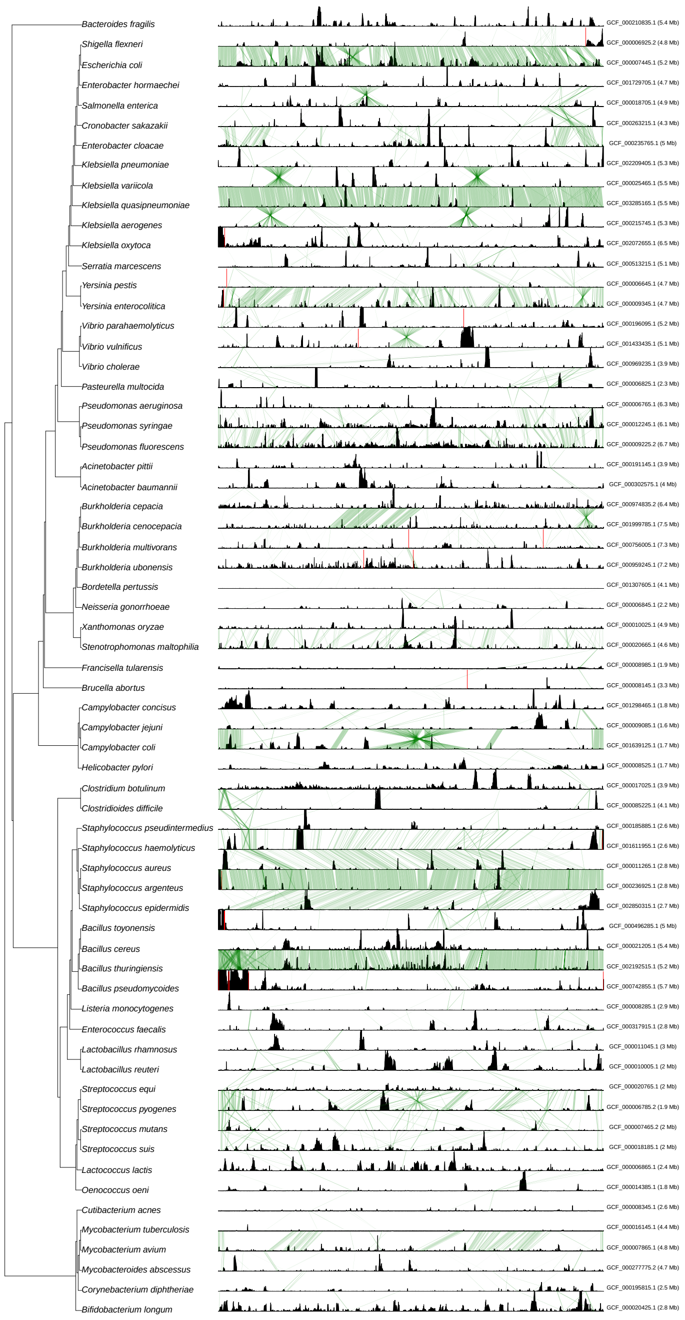


Figure 4: Comparison of complexity profiles of different bacterial species. Phylogenetic tree was build based on 16S rRNA sequence. Complexity profiles are shown in the same scale for all organisms. Synteny blocks are shown with green color. Genomes id used as reference are on the right. Red vertical lines separate replicons (plasmids or chromosomes). Only species with exactly 100 genomes used for complexity calculation are shown.

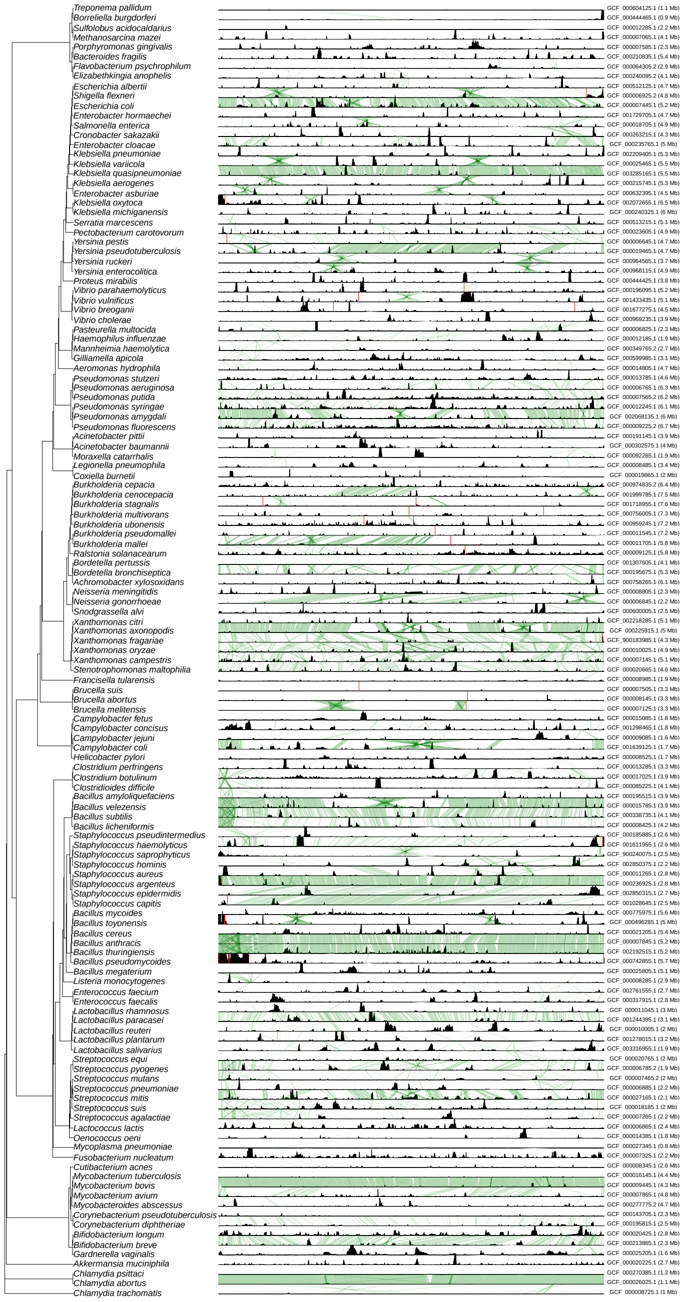


Figure 5: Comparison of complexity profiles of different bacterial species. Phylogenetic tree was build based on 16S rRNA sequence. Complexity profiles are shown in the same scale for all organisms. Synteny blocks are shown with green color. Genomes id used as reference are on the right.

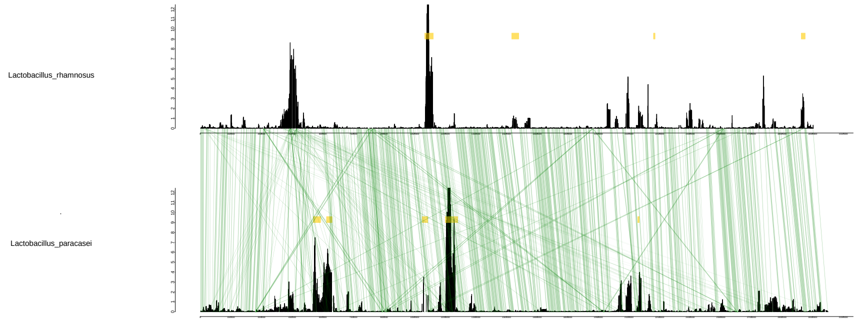


Figure 6: Comparison of complexity profiles of three species of Lactobacillus genus. Synteny blocks are shown with green color. Yellow rectangles denote prophage regions.

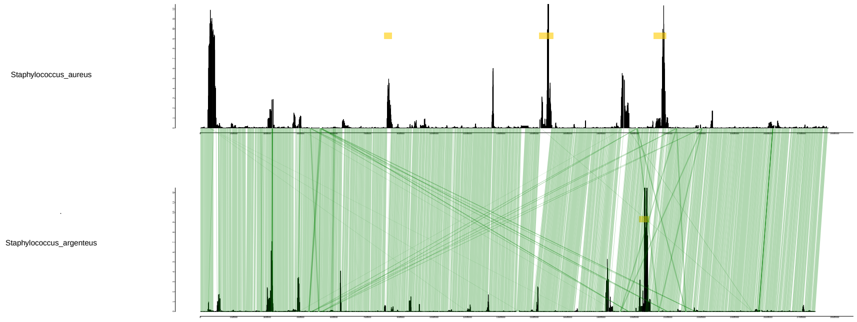


Figure 7: Comparison of complexity profiles of three species of Staphylococcus genus. Synteny blocks are shown with green color. Yellow rectangles denote prophage regions.

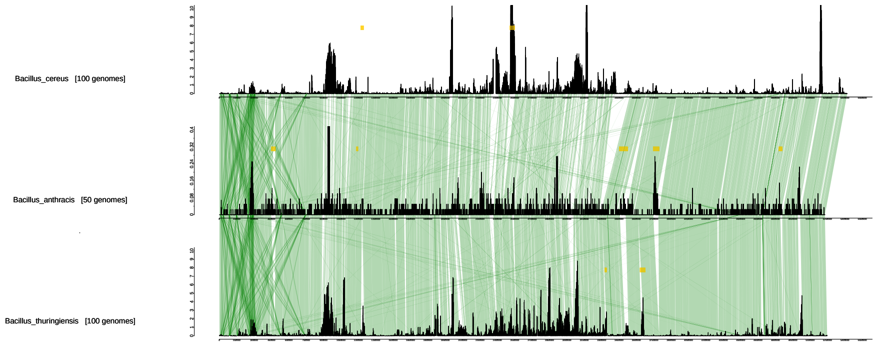


Figure 8: Comparison of complexity profiles of two species of Bacillus genues. Synteny blocks are shown with green color. Yellow rectangles denote prophage regions.

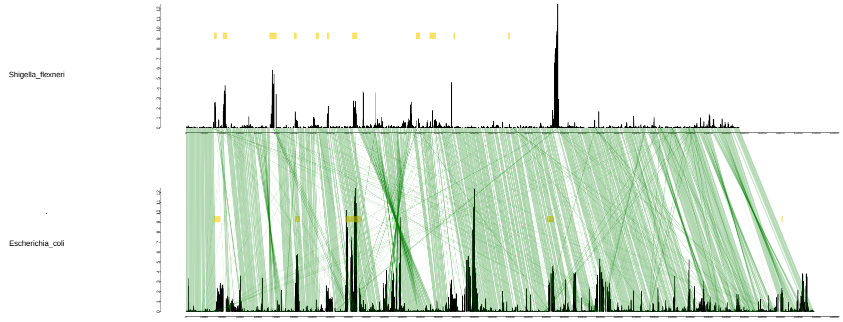


Figure 9: Comparison of complexity profiles of two genera of Enterobacteriaceae family. Synteny blocks are shown with green color. Yellow rectangles denote prophage regions.

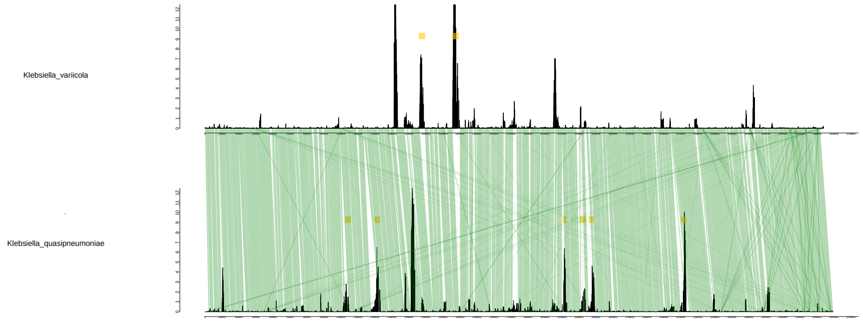


Figure 10: Comparison of complexity profiles of two species of Klebsiella genus. Synteny blocks are shown with green color. Yellow rectangles denote prophage regions.

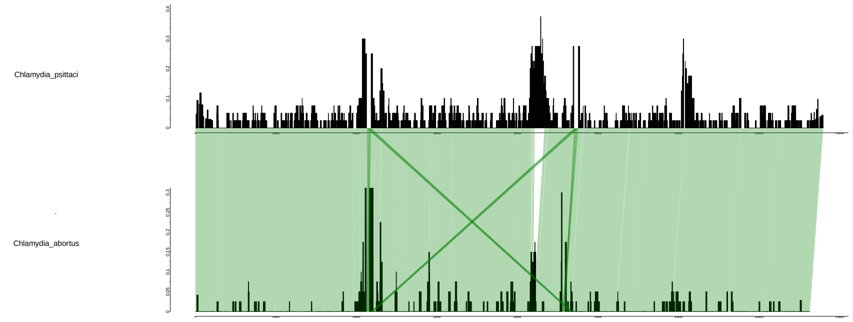


Figure 11: Comparison of complexity profiles of two species of Chlamidia genues. Synteny blocks are shown with green color. Yellow rectangles denote prophage regions.
