## Supplementary Notes for "Genome Complexity Browser: estimation and visualization of prokaryote genome variability"


1. **Paralogous problem.** **1**
2. **Alignment and graph building** **2**
3. **Genome complexity computing** **3**
4. **Subgraph drawing** **4**
5. **Genome rearrangements simulation.** **5**
6. **Libraries and frameworks** **7**
7. **Database structure** **7**
8. **Comparison of complexity values and nucleoid 3d conformation models.** **7**


### 1. Paralogous problem.

There is one problem in the graph building caused by the multicopy genes (paralogues), which can critically increase local graph connectivity and create loops. So, two methods were suggested to solve such situations.

The first method is a simple deletion of all orthology groups which have more than one copy in some genome from this genome. An advantage is simplicity and clear output. A disadvantage is a significant number of pseudodeletions (Figure 1A).

The second method does not delete paralogues, but “orthologizes” them (Figure 1B): paralogues groups are divided by their context. But this method creates a new problem with the appearance of many pseudoedges and pseudonodes that could increase the real value of genome complexity and noise in complexity distribution. There is one more process in the second method, is a collapsing of sequentially located identical nodes to one (for example, two *c* genes in G1 are collapsed to one *c_1* node).

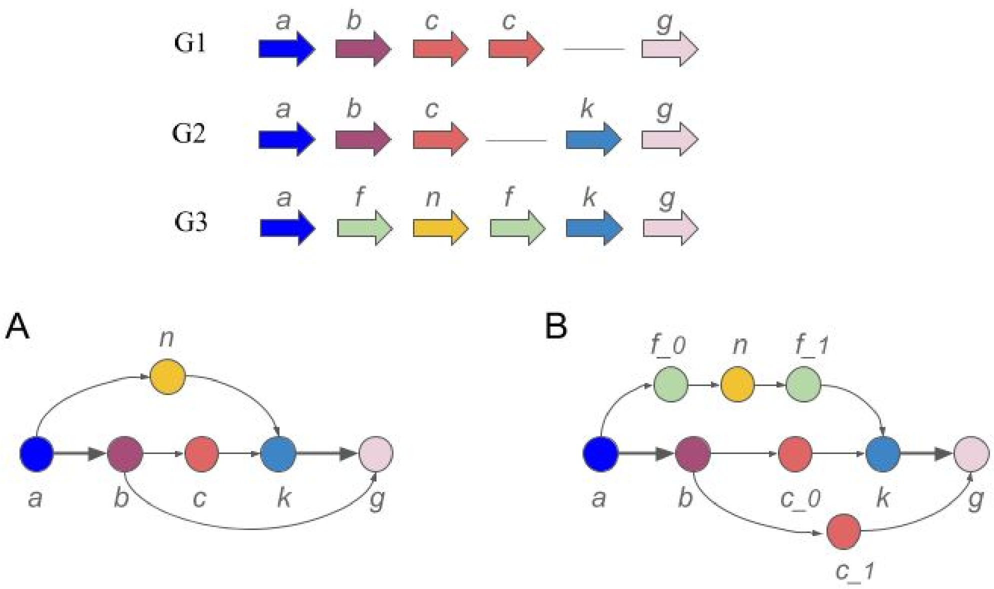


*Figure 1. Comparison of two methods to solve the paralogous problem;*
*(A) Simple paralogous deletion;*
*(B) Paralogous "orthologization".*

In GCB Web-based application both methods are available but we recommend to use the first to get a more clear output.

### 2. Alignment and graph building

Parsing and alignment of these orthology groups chains were done by a simple iterative algorithm. The first step is sorting all chains by their length. It is important to emphasize that one organism can contain more than one chain (for example in draft assemblies).

Next, the longest contig is selected as a reference. For each other contig in the set, it's checked that reference containing at least 50% of orthology groups from the current contig. If it is true, the orientations of all pairs of sequentially located orthology groups in reference and current contig are compared. If the count of reversed pairs orientation is more then count of non-reversed (considering the bp lengths of these pairs), we reverse the current contig. After considering all contigs a set of unprocessed contigs can remain because of the 50% filter. If it is true, the longest contig from these non-compared set is selected as a new reference contig and the procedure is repeated. The cycle repeats until all contigs will be compared.

Finally, all compared contigs are written to a file (default *paths.sif*). This file content aligned orthology group sequences for each sampled organism and used in all graph processing scripts. Database using in GCB service is generated simultaneously.

This alignment algorithm has the disadvantage that not all contig are aligned with each other, which can cause the appearance of extra edges in the graph structure. This problem can be partially solved by reducing of the 50% filter, but too small value of this filer can lead to reversing error because a difference between the count of reversed and non-reversed pairs orientations will become less significant.

Next, the graph structure will be created using *paths.sif* file.

All orthology groups are defined as nodes of the graph. The directed edges are conducted between nodes which are located sequentially in at least one genome of the sampled organisms.

For simple and fast processing the graph is structured as a hash table (an ordered dictionary in the Python implementation), where each node is the key, and the set of all nodes to which the edges from the key-node are drawn is stored value. To decrease memory usage all nodes coded as integer numbers.

### 3. Genome complexity computing

First, it is necessary to choose the reference organism. Complexity in each node (orthology group) which belong to the genome of the reference organism will be computed iteratively.

We define the graph complexity in a node as the number of possible unique deviating paths in the graph that begin and end within the user-defined window. Deviating paths are defined as graph paths that begin and end in the reference sequence of nodes, but differ from the reference at all other nodes (Figure 2).

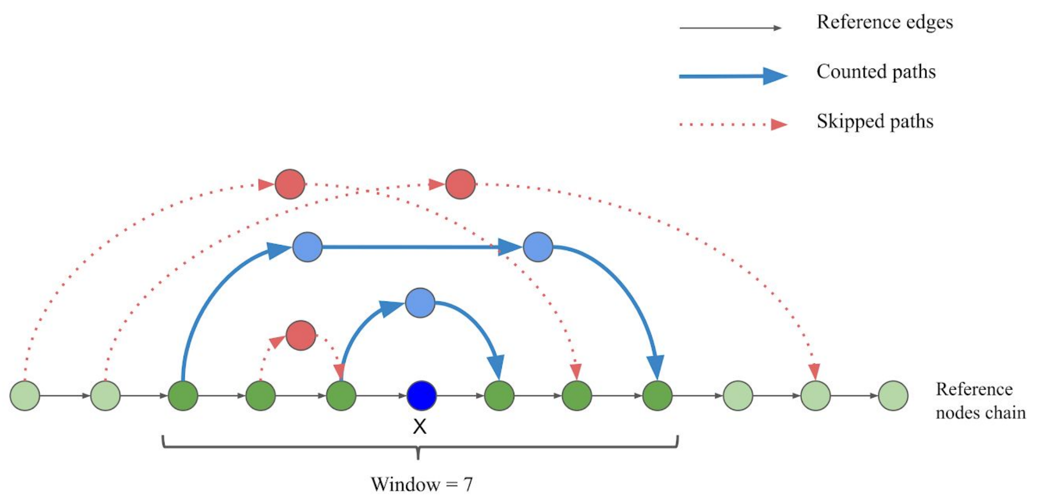


*Figure 2. Counted deviating paths are drawn by blue color. Skipped paths are not counted as deviating paths because of the user-defined window parameter.*

To search deviating paths a probabilistic algorithm is proposed. It is based on a random walk through the graph and takes as an argument the number of iterations of searching for random paths from each node of the referency chain. A random walk was chosen as a substitution for a full search, which becomes computationally complicated on a sufficiently large graph.

Before the deviating paths search, the complexity of each node is assumed to be zero. After searching for each unique deviating path for all nodes lying between the output and the input of this path, the complexity value increases by window normalized value. Thus, after searching for all deviating paths in the graph, we have a table of complexity values for all nodes from the reference chain.

### 4. Subgraph drawing

Our tool allows to draw a graph and describe complexity in any graph region but visualization of the full graph is too difficult to understand and execute. So, it was being decided to visualize only a small part of the graph, which consists only of the context of some gene or group of genes.
To construct a subgraph consisting only of nodes and edges lying on the region of interest, we proposed an iterative algorithm (Supplementary Listings 1, LISTING 6).
In GCB service we use subgraph structure to draw any interest region and its context. To render a subgraph image we use Cytoscape.js library, which allows users to manipulate with a rendered graph: move nodes, select nodes sets, etc. In our opinion, the most useful and clear render layouts are produced by Dagre and Graphviz.
In the Web-based application, the rendering stage has few user-defined parameters. Part of them is a set of subgraph generating parameters, such as *window* (default 20), *tails* (default 5), *depth* (default 30), *reference genome* and *coordinates* of interest region. Other options control only the rendering process. *Minimal edge weight* is a numeric parameter filtering edges which belong to less than this number of genomes. *Layouter* allows changing layout rendering algorithm.
Drawed edges weight is proportional to the square root of the number of genomes and calculated as

$$10 * \sqrt{\frac{n}{max(n)}}$$

to provide normalized weights for sets with any number of genomes.

### 5. Genome rearrangements simulation.

To validate our method we assumed non-random distribution of the probability of genome rearrangement events in different genome locations. First, we created 100 identical genomes and then run random genome rearrangements simulations with 3000 iterations. We used three predefined patterns (sinus, sawtooth, rectangular) of the probability distribution (Figure 3) through the genome.

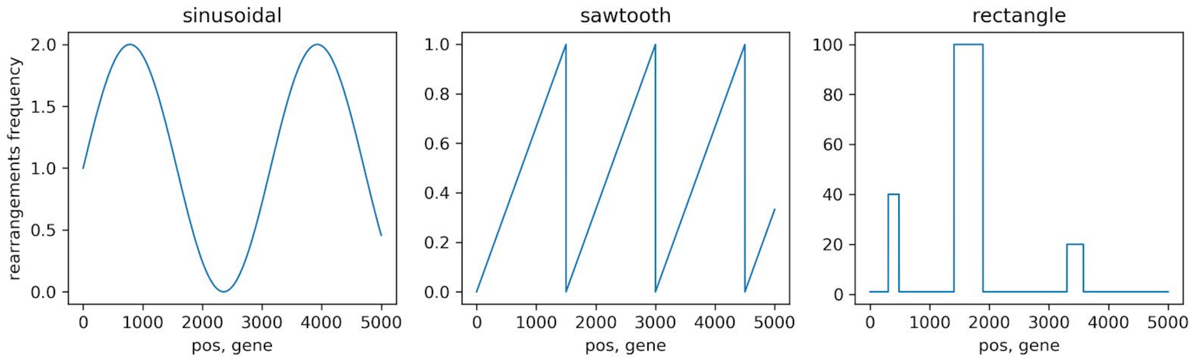


*Figure 3. Pre-defined distribution: sinusoidal, sawtooth, rectangular*

For each iteration, we performed genome changes: insertion of a new gene(s) (from "orbital" genes set), horizontal gene transfer (HGT) from other genomes, deletion of the gene(s), inversion of part of the genome. The total number of genes in any genome was constant.
Length of each rearrangement was determined by an exponential distribution with $1/\lambda = 0.005$.
For each distribution type, 10 independent simulations were performed.
After all iterations, generated genomes were used to generate gene context graph and our algorithm of complexity estimation was applied and compared to the initial rearrangement distribution (Figure 4).

*Simulations parameters:*
insertion probability = 0.5
HGT probability = 0.5
inversion probability = 0.005
genome length = 5000
number of genomes = 100
number of iterations = 3000

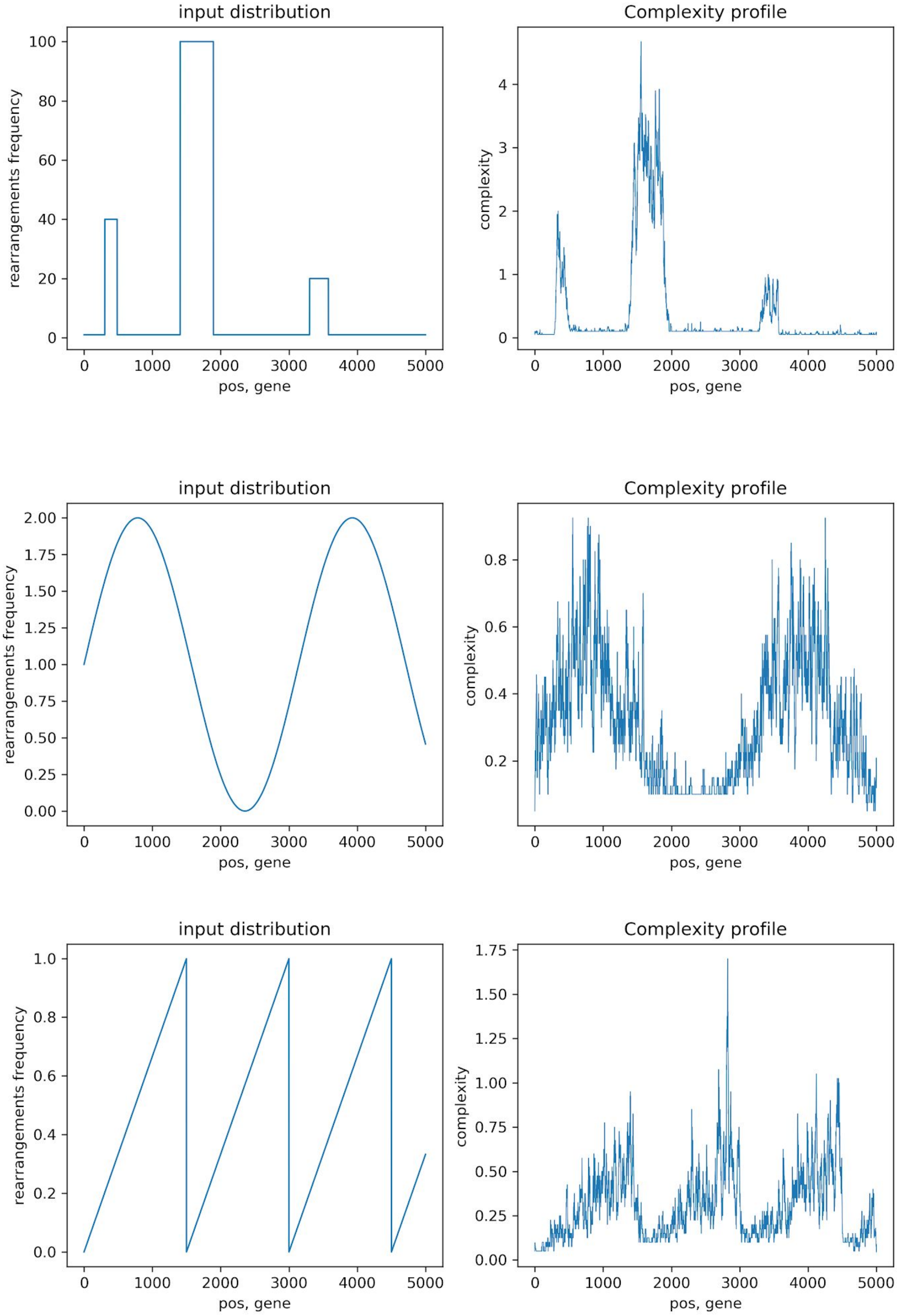


*Figure 4. Complexity computing results for simulated datasets.*

To estimate the similarity between input and output distributions R-square value and Spearman correlation coefficient was used (Table 1). FDR Benjamini-Hochberg multitest correction was applied to correct p-values of these tests for each model. In all cases calculated p-values were smaller than floating-point type numbers threshold ($< 10^{-308}$).

Table 1

**Estimation of the method quality**

| Input distribution | R-square mean | Spearman coefficient mean |
| --- | --- | --- |
| sinusoidal | 0.767 | 0.822 |
| rectangle | 0.946 | 0.699 |
| sawtooth | 0.800 | 0.847 |

It is possible to carry out the validation pipeline with one sinusoidal distribution by a script that is available on http://github.com//DNKonanov/geneGraph/. Validation script “validate.sh” is located in “geneGraph-master/source/validation/”. This short pipeline includes random rearrangements simulations, complexity computing and results plotting.

### 6. Libraries and frameworks

We use React+Redux to render application page and store client-viewed data. Requests are got and processed by Flask web-server framework. Complexity plots are produced by Plotly.js library. SQLite databases are used to store all service data.

### 7. Database structure

SQLite database (DB) is used to store GCB service data. For each organism, DB file is created separately to optimize data access and searching and simplify new organisms adding. DB consists of six tables:

1) Genomes table - stores information about genomes, including RefSeq codes, real strain names, and ids.
2) Contigs table - stores information about all contigs, them codes, ids and genomes to which these contigs belong.
3) Nodes table - stores nodes (genes) sequences for each contig, them coordinates, and annotations.
4) Edges table - stores information about all edges in the graph, them weights and genomes to which these edges belong.
5) Complexity table - stores complexity profiles for all genomes.
6) Node keys table - stores information about orthogroups names coding, which is used to decrease memory usage in graph processing.

### 8. Comparison of complexity values and nucleoid 3d conformation models.

We supposed that the rearrangements frequency of some region can depend on the location of this region in the chromosome globule. For example, genes located on the edge of the globule may be of different complexity than genes located in the globule center. To check this assumption we used 3D nucleoid models generated by the plectoneme-based method developed by (Hacker and Elcock, 2017)

To observe relation between gene complexity and gene location in the globule we approximated nucleoid as the cylinder with semi-spherical bases. Next, we divided the volume of this cylinder on two parts: outer and inner by another cylinder, in such a way that inner and outer volumes contain the same or almost the same number of genes (Figure 5). Both oriC@pole and oriC@midcell models were used in the analysis, 10 structures for each model.

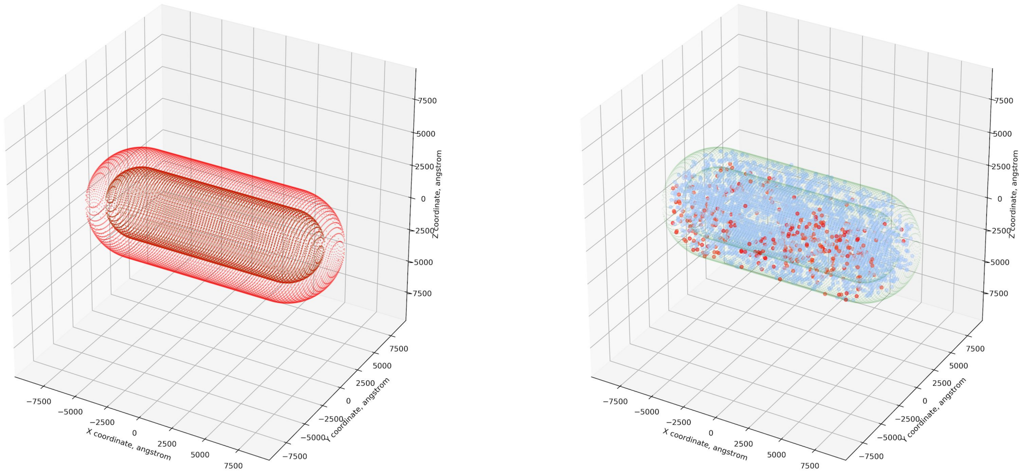


A B

*Figure 5. The left plot (A) shows the cylinders with semi-spherical bases modeling chromosome globule (outer and inner parts). The right plot (B) shows the location of all genes plotted as points in the globule model. Genes depicted by red points have greater than half of the maximum complexity; orange points have complexity between half and lower quartile; blue - lower than lower quartile.*

Mann-Whitney test was used to estimate the difference between the complexity of inner and outer genes. There were no statistically significant results in this comparison (FDR corrected p-values > 0.3).

**Comparison of complexity values and intrachromosomal interactions data.**

One of the outputs of GCB is a table of numbers of unique deviating paths between all possible pairs of nodes (genes) (may be called bridges or arcs). These numbers can be considered as a rearrangements frequency between these genome positions. We performed a comparison of these numbers with experimentally observed intrachromosomal interactions. Prokaryotic chromosomes, as well as eukaryotic, are known to have domain structure (Postov et al., 2004), so, we supposed that a similar domain structure may also appear in directions and localizations of genome rearrangements.

We used 3C data computed for *Escherichia coli MG1655* (Lioy et al., 2018). All bridges calculated for this organism by our method were represented as a heatmap of rearrangements density between all genome positions (Figure 6).

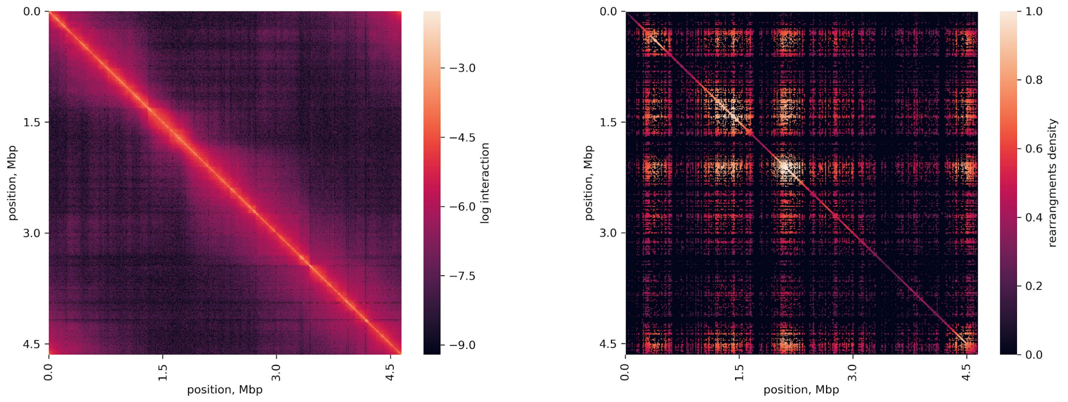


*Figure 6. Comparison of 3C data and density of deviating paths between all genes positions.*

It can be seen that there is a number of domain structures in both 3C and rearrangements density heatmaps but we did not find statistically significant evidence that there is a similarity between them. Matrices were compared by Mentel test, correlation coefficient = 0.036 (Spearman rank test to compute correlation was used).

Comparison of cumulative 3C values that "reflect the relative tightness of the distribution of contacts made by the chromosomal region" (Lioy et al., 2018) - with complexity values of *E. coli MG1655* did not reveal any apparent regularities (Figure 7).

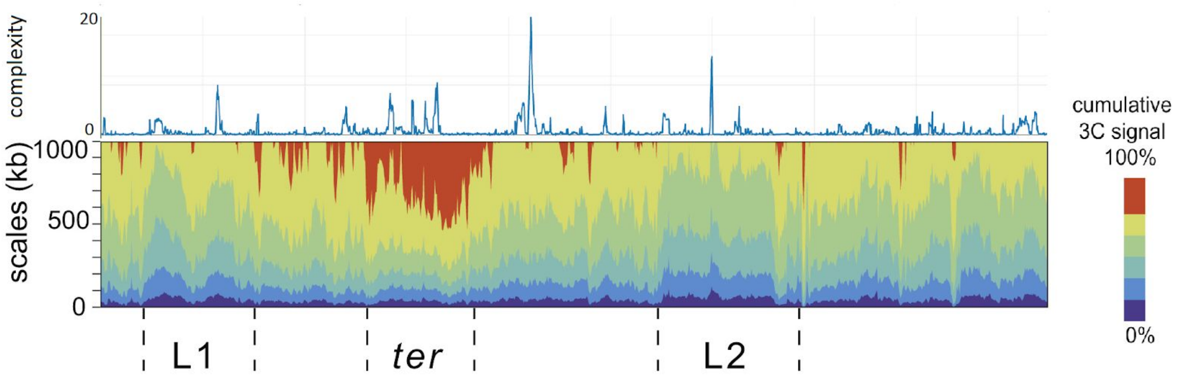


*Figure 7. Comparison of 3C data presented as cumulative values with complexity values. Scalogram image is from (Lioy et al., 2018).*
