## Supplementary Listings for "Genome Complexity Browser: estimation and visualization of prokaryote genome variability"


LISTING 1. ALIGN GENOMES SET 1

LISTING 2. ORTHOLOGIZE PARALOGUES GROUPS 2

LISTING 3. FIND PATHS 2

LISTING 4. COMPUTE COMPLEXITY 3

LISTING 5. GENOME REARRANGEMENTS SIMULATION 3

LISTING 6. GENERATE SUBGRAPH 4


---

### LISTING 1. ALIGN GENOMES SET

Input: non-reversed nodes chains
Output: list of aligned genomes

all_contigs ← get full contigs list from non-reversed chains
sort all_contigs by their length

aligned_contigs ← empty list

**while** all_contig is **not** empty **do**

    reference_contig ← longest (first) contig from all_contigs

    **for each** contig **in** all_contigs **do**
        **if** contig is reference_contig **do**
            move contig from all_contig to aligned_contigs

        **if** reference_contig do not contains at least 50% of nodes from contig **do**
            **continue**

        forward_count ← count of pair of sequently located gene contained in both reference_contig and contig
        reverse_count ← count of pair of sequently located gene contained in both reversed reference_contig and contig

        //forward_count and reversed_count calculated with multiplication by the length of the corresponding genes

        **if** reverese_count > forward_count **do**
            move reversed contig from all_contigs to aligned_contigs

        **else do**
            move contig from all_contigs to aligned_contig

list of aligned genomes ←sort aligned contigs by organisms
**return** list of aligned genomes

---

### LISTING 2. ORTHOLOGIZE PARALOGUES GROUPS

Input: list containing nodes chains (contigs) with paralogues for each genome
Output: genomes with “orthologized” paralogous

paralogues_set ← list of nodes, which have more than one copy in at least one genome
contexts_set ← dictionary containing list of all possible contexts for each paralogues, initialized as empty

```
for each genome in input list do
        for each contig in genome do
                i = 1
                while (i < length(contig) - 1) do
                        if contig[i] not in paralogues_set do
                                i = i+1
                                continue
                        if contig[i-1] == contig[i] do
                                merge contig[i] and contig[i+1] to contig[i]
                                continue
                        context = (contig[i-1], contig[i+1])
                        if context not in contexts_set[contig[i]] do
                                add context to contexts_set[contig[i]]
                                rename contig[i] to new unique node and bind this name with this context
                        else if context in contexts_set[contig[i]] do
                                rename contig[i] to existing name bound with this context
                        i = i+1

return modified genomes list
```

### LISTING 3. FIND PATHS

// graph is stored as a hash table, where each node is key and each value is set of nodes connected by an edge with key-node

Input: graph, start_node, base line, number of iterations
Output: Paths - set of all paths, which start in start_node and ends in base line

```
Paths ← empty set
i ← 1
while (1 <= i <= iterations) do
        path = [start_node]
        next_node ← select random node connected with current_node
        while next_node not in base_line do
                if next_node not in path do // to avoid cycle paths
                        add next_node to path
                next_node ← select random node connected with the last node in the path
        if length(path) > 1 and path not in Paths do
                add path to Paths
        i ← i + 1
return Paths
```

### LISTING 4. COMPUTE COMPLEXITY

```
Input: graph, reference organism, window size, number of iterations
Output: complexity values for each node in the reference
ref_chain ← reference nodes chain
start complexity value for all nodes ← 0
for each node in ref_chain do
        Paths = FIND PATHS (node) // find all paths that start in node and end in ref_chain
        for each path in Paths do
                start ← first node in the path
                end ← last node in the path
                distance ← | position(start) - position(end) |
                if (distance <= window) do
                        for each node between start and end do
                                complexity[node] ← complexity[node] + 1/(2*window)
return complexity values
```

### LISTING 5. GENOME REARRANGEMENTS SIMULATION

```
Input: changes frequency distribution, probability of different changes (HGT, deletion, inversion), exponential parameters of changes length, number of genomes, number of iterations, number of "orbital" genes
Output: graph.sif file with simulated genomes set

genomes ← set of identical genomes with 5000 genes
i ← 0
while i < number of iterations do
        genome ← random genome from genomes
        //random(0,1) generates random float number between 0 and 1
        if random(0,1) <= inversion_probability do
                inversion_length ← random value from exponential distribution with 1/λ = input exponential parameter
                inversion_position ← randomly generated position in genome based on input distribution
                apply inversion

        if random(0,1) <= insertion_probability do
                insertion_length ← random value from exponential distribution with 1/λ = input exponential parameter
                insertion_position ← randomly generated position in genome based on input distribution
                inserton_chain ← random list of "orbital" genes with length insertion_length
                // "orbital" genes is set of genes named from 5000 to 5000+number of "orbital" genes
                apply insertion
                apply random deletion //It's necessary because we need to store the same number genes in genomes

        if random(0,1) <= HGT_probability do
                other_genome genome ← random other genome from genomes
                HT_length ← random value from exponential distribution with 1/λ = input exponential parameter
                HT_position_from ← randomly generated position in genome based on input distribution
                HT_position_to ← randomly generated position in other_genome based on input distribution
                apply HT
                apply random deletion from other_genome
        i ← i + 1
write genomes to *.sif file
// organism are named as org0, org1, …, orgN and org_ref, where org_ref is no-changed chain (from 0 to 4999)
// NB!: only one copy of each gene in one genome are allowed
return
```

### LISTING 6. GENERATE SUBGRAPH

```
Input: graph, reference, start_node, end_node, window, depth, tails, minimal_edge_weight
Output: subgraph

subgraph ← empty graph

target_chain ←nodes chain from reference genome between start_node - window and end_node + window
add base_chain to subgraph

deviating_paths ← find all deviating paths connected with target_chain

for each path in deviating_paths do
        if length of the path <= depth do
                add path to the subgraph
        else do
                path_tails ←left and right fragments of the path with length tails
                add path_tails to the subgraph

for each edge in subgraph do
        if edge weight < minimal_edge_weght do
                delete edge from the subgraph

clear the subgraph from disconnected fields

return subgraph
```
